# Inferring somatic copy number aberrations from spatial transcriptomics data to study early cancer development

**DOI:** 10.64898/2026.10.23.761433

**Authors:** F Honig, C Brochen, M Basha, R Arora, M Jimenez-Linan, AC Antoniou, OM Rueda, F Correia Martins

## Abstract

Understanding the spatial organisation of cells undergoing early transformation and their interactions with the local microenvironment is critical to studying the earliest stages of cancer development. Somatic copy number aberrations (SCNAs) are among the earliest genomic alterations observed in many cancers, but their spatial context remains difficult to study using conventional sequencing approaches. We developed a computational framework to infer SCNAs directly from Xenium spatial transcriptomics data, enabling the identification and spatial mapping of cell populations sharing common SCNA profiles. Applying this approach to ovarian, cervical and breast cancer datasets identified spatially coherent SCNA-defined populations that recapitulated known tumour-associated copy number aberrations and pathological states. In high-grade serous ovarian cancer, we identified an intermediate epithelial population within the fallopian tube exhibiting features of early transformation accompanied by immune remodelling. Together, our pipeline establishes a scalable framework for reconstructing the spatial architecture of SCNA-defined clones, providing new opportunities to identify and characterise candidate early transforming cell populations in situ.

## Introduction

Chromosomal instability (CIN) is an ongoing process that generates somatic chromosomal copy number alterations (SCNAs), potentially disrupting cancer-relevant genes and shaping tumour initiation, progression and response to therapies^1–3^. High-grade serous ovarian cancer (HGSOC) and triple-negative breast cancers are characterised by extensive SCNAs, with early CIN and *TP53* mutations detectable in precursor lesions in the fallopian tubes (including secretory epithelial cells - SECs) and breast, respectively^4–8^.

Aneuploidy is uncommon in normal tissues in the presence of functional wild-type *TP53*, whereas patches of *TP53*-mutant cells are widespread across normal tissues^9–11^. Moreover, most carriers of germline *TP53* mutations do not develop cancer in every tissue^12,13^. Together these observations indicate that *TP53* mutation alone is insufficient to drive cancer transformation and may instead create a permissive context in which additional genomic alterations, including SCNAs, accumulate. CIN and associated replication stress can trigger cell cycle arrest and apoptosis^14–16^, while chromosomal missegregation can generate micronuclei that activate cGAS and innate immune signalling^17,18^. CIN may therefore influence not only the survival of genomically unstable cells but also their interactions with surrounding stromal and immune cells.

However, how emerging genetically altered clones interact with their tissue environment during the earliest stages of transformation remains poorly understood. Addressing this question requires simultaneous resolution of genomic alterations, cellular states and spatial organisation. Previous approaches have integrated single-cell DNA sequencing and transcriptomic data to associate SCNAs with cellular phenotypes, exploiting the SCNA-induced dosage effect on gene expression^19–21^. Other approaches have improved SCNA inference by exploiting allele-specific effects^22–24^. Despite linking genomic alterations with cell states, these advances provide limited information about the spatial organisation of genetically defined clones within tissue ecosystems.

Recent improvements in spatial transcriptomics provide an opportunity to address this gap. Xenium 5K enables highly multiplexed, spatially resolved single-cell transcriptomic profiling^25^ . By integrating gene-dosage effects across multiple genes with the spatial proximity of genetically related cells, spatial transcriptomic data can potentially be used to infer SCNAs and reconstruct the organisation of genetically defined clones in situ^26^.

Here, we develop a computational framework to infer SCNAs from Xenium spatial transcriptomic data and identify genetically defined cell clones. Beyond spatial SCNA inference and clone detection, the framework facilitates a broad range of downstream analyses, including characterisation of SCNA-defined transcriptional programmes through differential expression analysis, gene ontology enrichment analysis, cell type composition profiling, identification of local genomic density hotspots and key gene clusters, and spatial mapping of SCNA-defined cell populations within their tissue microenvironment. We apply this approach to histologically normal tissues to detect early genetic alterations that may be missed by conventional pathological assessment, and to define their cellular and spatial context. In the fallopian tube, the site of origin of HGSOC, we use this framework to investigate how emerging abnormal clones interact with the immune microenvironment, providing a spatial view of immune surveillance and potential immune evasion during the earliest stages of malignant transformation.

## Results

### Chromosome arm-level expression enables SCNA inference using diploid reference

To determine whether gene-dosage effects captured by spatial transcriptomics could be used to infer SCNAs, we analysed a publicly available Xenium 5K human ovarian cancer dataset (10x Genomics), containing epithelial, stromal and immune cell populations (Figure 1A). We reasoned that broad SCNAs should produce coordinated changes in the expression of multiple genes located within the same chromosome arm, allowing copy number states to be distinguished from gene-specific transcriptional variation.

**Figure 1.**
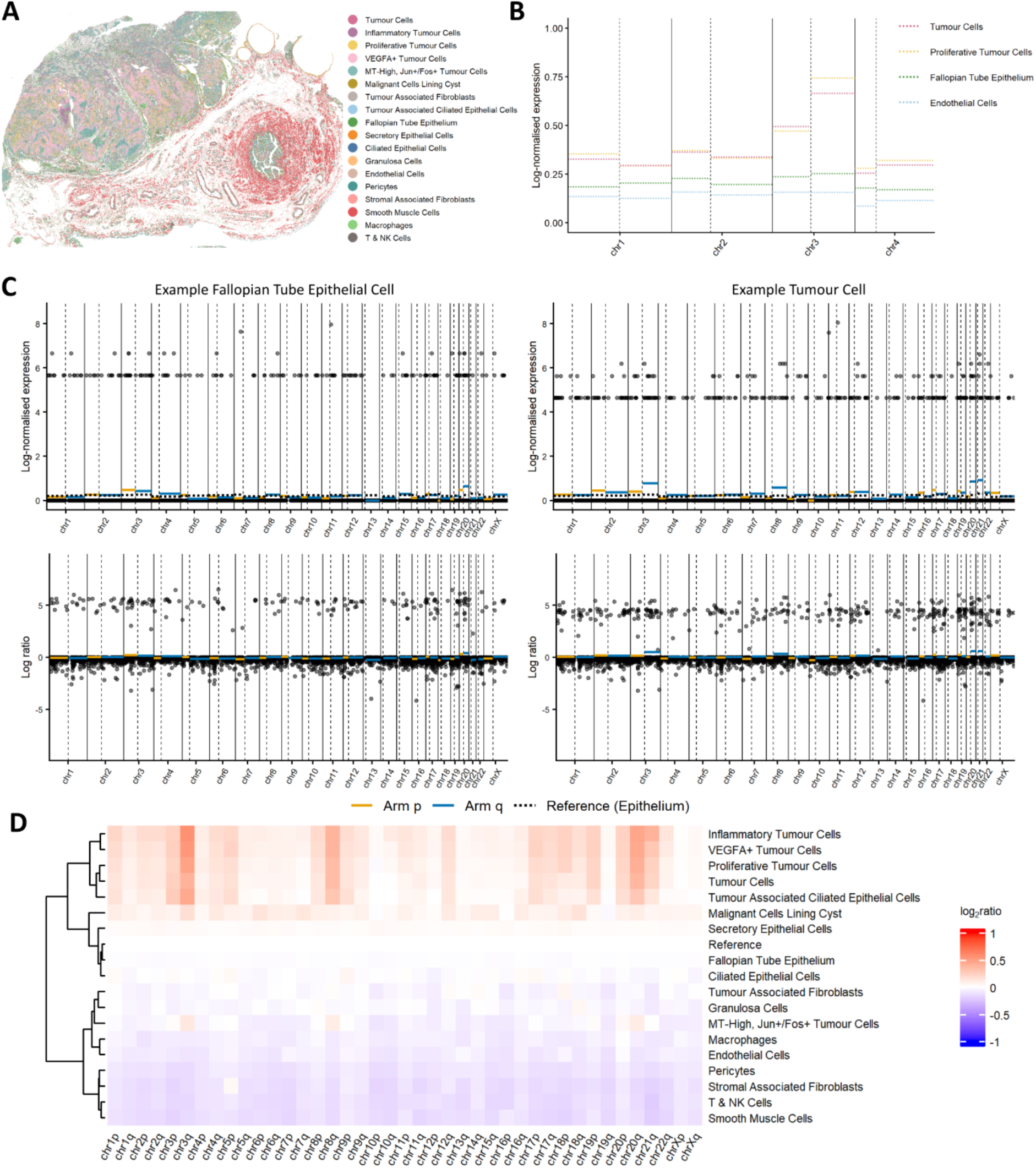
Chromosome arm-level gene expression aggregation identifies somatic copy number aberrations (SCNA) in tumour cell populations. **(A)** Spatial distribution of annotated cell populations in Xenium 5K human ovarian cancer spatial transcriptomics dataset (10x Genomics). Colours indicate cell type. **(B)** Mean log-normalised chromosome arm-level gene expression across representative cell populations. **(C)** Representative fallopian tube epithelial (left) and tumour (right) cells showing gene-level and arm-level expression profiles (top: short arm, p, orange; long arm, q, blue) and SCNA log ratios relative to the mean epithelium diploid reference (bottom: green dashed line). **(D)** Heatmap of chromosome arm-level SCNA log ratios across annotated cell populations.

Following log2-normalisation, we therefore aggregated expression across genes within each chromosome arm to generate chromosome arm-level expression profiles for individual cells. Comparison of these profiles across annotated cell populations revealed marked differences between non-malignant and tumour-associated cell populations (Figure 1B; Supplementary Figure 1). Whereas non-malignant epithelial, stromal and immune populations showed relatively uniform chromosome arm-level expression, tumour cells exhibited recurrent coordinated deviations affecting multiple chromosome arms.

We next established a diploid reference using the mean chromosome arm-level expression profile of histologically normal fallopian tube epithelial cells, including SECs. Comparison of individual cells to this reference revealed that the chromosome arm-level expression of a representative fallopian tube epithelial cell closely matched the diploid baseline, whereas a tumour cell showed pronounced deviations across multiple chromosome arms, including increased relative expression of genes on chr3q and chr8q (Figure 1C; top).

To convert these expression differences into inferred SCNAs, we computed the deviation in chromosomal arm-level expression relative to the diploid reference for each cell, generating log ratio profiles (Figure 1C; bottom). Applying this approach across all cells, the inferred SCNA profiles by annotated cell type identified recurrent chromosome arm-level SCNAs in tumour-associated cell populations, whereas non-malignant populations remained largely copy number neutral (Figure 1D). Altogether, these results support the use of chromosome arm-level expression as a basis for inferring broad SCNAs from Xenium 5K spatial transcriptomics data, distinguishing genomically abnormal tumour-associated populations from non-malignant cells.

### Spatial clustering resolves distinct SCNA populations

We next incorporated spatial information using the previously published BANKSY algorithm^27^. It was applied to the inferred chromosome arm-level SCNA log ratios, integrating neighbourhood composition and local spatial structure to identify spatially coherent cell populations with shared SCNA profiles. Optimisation of the mixing parameter (*λ*) and clustering resolution (res) enabled stable cluster identification, with *λ* = 0.8 and res = 0.1 providing the clearest separation of biologically distinct populations (Supplementary Figure 2). This approach resolved four major spatial clusters characterised by distinct chromosome arm-level SCNA profiles (Figure 2A).

**Figure 2.**
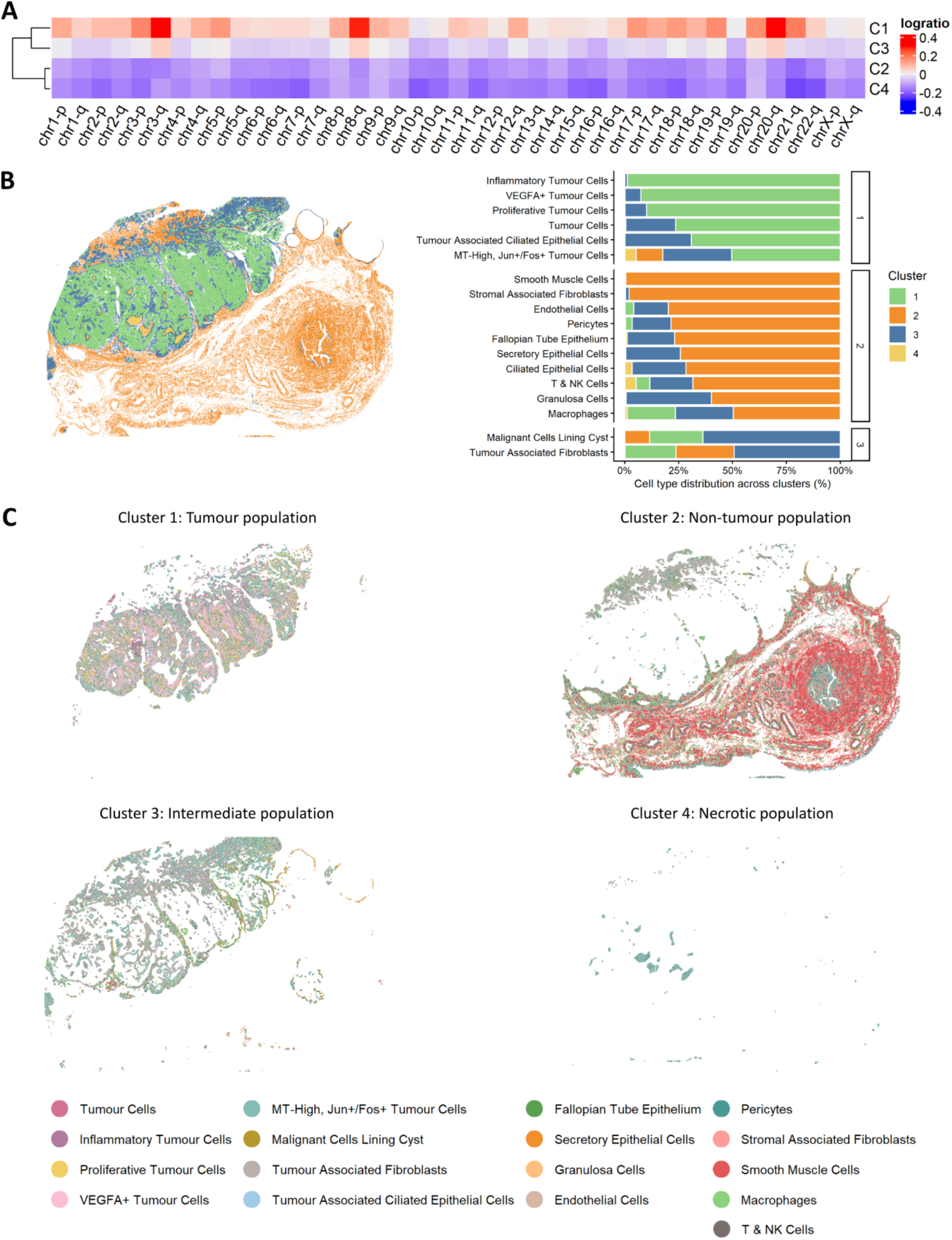
Spatial clustering of chromosomal SCNAs reveals spatial cell populations reflecting distinct stages of evolutionary cancer progression. **(A)** Heatmap of spatially inferred chromosome arm-level SCNAs. **(B)** Spatial distribution of SCNA-defined clusters across the tissue section (left) and corresponding cell type distribution across each cluster (right). Cell types are grouped and ranked by prevalence across clusters. Colours indicate spatial cluster. **(C)** Spatial distribution of individual clusters annotated by cell type, categorised as tumour (Cluster 1), non-tumour (Cluster 2), intermediate (Cluster 3) and necrotic (Cluster 4) cell populations. Colours indicate cell types.

Cluster 1 displayed widespread inferred gains across multiple chromosome arms, including 3q, 8q and 20q, whereas Cluster 2 was largely copy number neutral. Interestingly, Cluster 3 showed an intermediate profile, with more modest gains affecting several of the same chromosome arms, while Cluster 4 displayed a distinct but largely neutral copy number profile. Spatial mapping demonstrated that Cluster 1 localised predominantly to the tumour region, Cluster 2 to normal epithelial and stromal cell populations, and Cluster 3 mainly adjacent to the tumour region, with a subset of cells within histologically normal fallopian tube epithelium (Figure 2B; left). Cluster 4 occupied tissue regions that were independently annotated as necrotic by pathological assessment. Analysis of the cellular composition of each cluster, ranked by prevalence across clusters, further revealed differences in cell type abundance (Figure 2B; right). Tumour-associated cell types were most abundant in Cluster 1, whereas epithelial, stromal and immune cell populations dominated Cluster 2. Consistent with its intermediate SCNA profile, Cluster 3 was enriched for malignant cells and tumour-associated fibroblasts, while maintaining representation from most other cell types. Based on their SCNA profiles, spatial localisation and cellular composition, the four clusters were subsequently annotated as tumour (Cluster 1), non-tumour (Cluster 2), intermediate (Cluster 3) and necrotic (Cluster 4) populations for downstream analyses (Figure 2C).

To assess biological plausibility, we compared the inferred SCNAs with the genome-wide SCNA frequencies from the TCGA Ovarian Serous Cystadenocarcinoma PanCancer Atlas^4,28,29^. Recurrent gains and amplifications were most prominent on chromosome arms 3q and 8q (Supplementary Figure 3), closely mirroring the dominant SCNA gains observed in the tumour population (Cluster 1). This agreement with the TCGA data provides orthogonal support for the biological relevance of the inferred spatial SCNAs.

### Genomic clustering of genes underlying SCNA populations

We next asked whether genes driving the expression-based SCNA signal localised to genomic regions implicated in HGSOC. Differential expression between tumour (Cluster 1) and non-tumour (Cluster 2) populations identified 529 upregulated genes represented on the Xenium 5K panel. Their genomic distribution was non-uniform (Figure 3A), with local density peaks overlapping regions containing established HGSOC drivers^3,4,30,31^, including *MECOM*, *PIK3CA*, *MYC* and *CCND1*, and coincided with the most frequently amplified regions in TCGA HGSOC cohort^4,28,29^ (Supplementary Figure 3).

**Figure 3.**
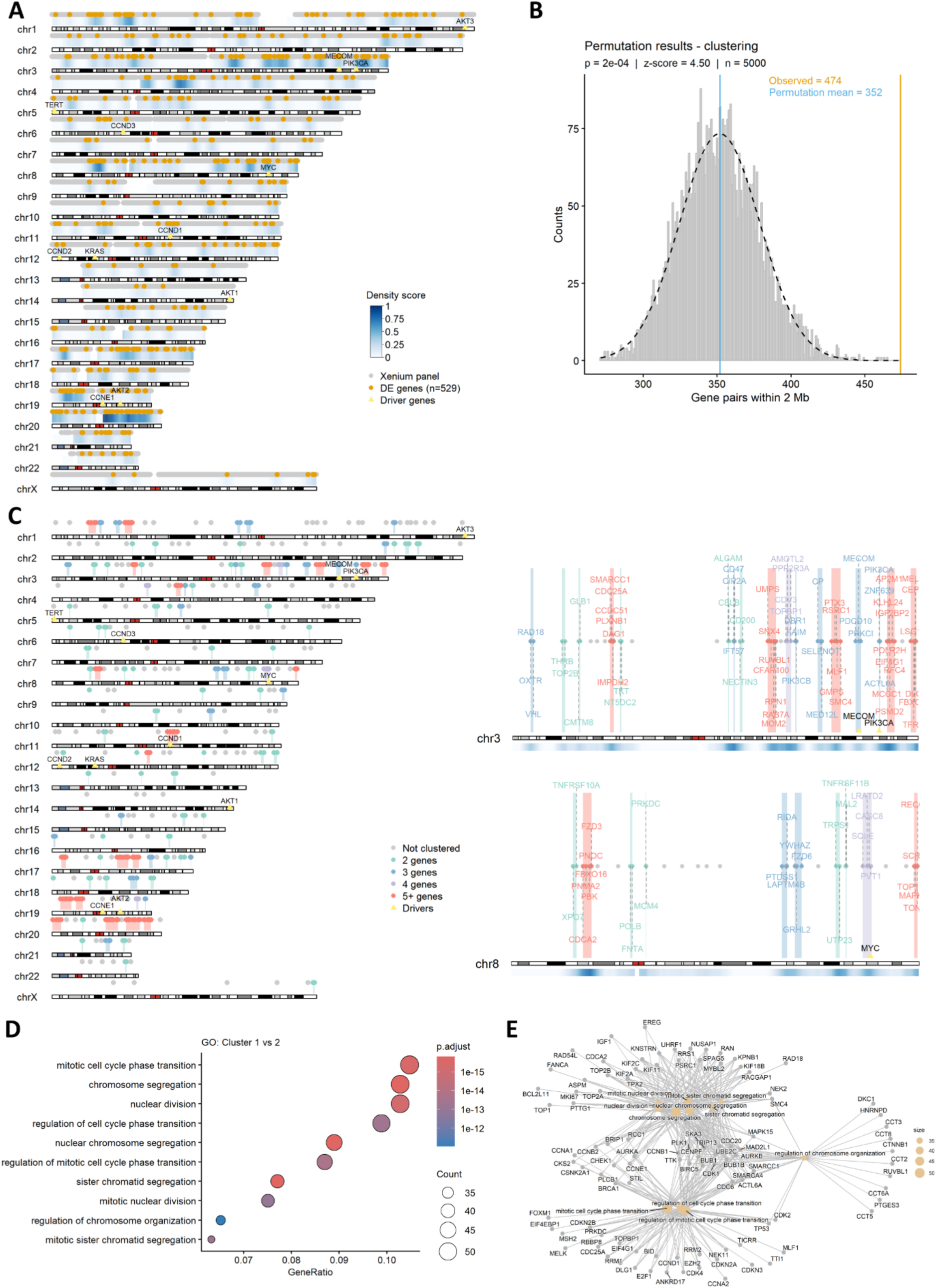
Key genes underlying spatial SCNA populations form local genomic clusters and are associated with chromosomal instability. **(A)** Chromosome track visualisation of the genomic distribution of upregulated genes (orange) in the tumour population (Cluster 1) relative to the non-tumour population (Cluster 2), shown against the complete Xenium 5K panel (grey). Heatmap indicates the local density of upregulated genes across the genome. Known cancer drivers are indicated by yellow triangles. **(B)** Permutation analysis assessing genomic clustering of the upregulated genes shown in **(A)** by quantifying gene pairs located within 2 Mb of one another following random gene shuffling (n = 5000). **(C)** Genomic distribution of clusters of neighbouring genes located within 2 Mb of one another, grouped by cluster size (2 = green, 3 = blue, 4 = purple, ≥ 5 = red). Insets show representative high-density gene clusters on chromosomes 3 and 8 associated with the most pronounced SCNAs. Known cancer drivers are indicated by yellow triangles. Local gene density is shown below each genomic track. **(D)** GO enrichment analysis of genes upregulated in tumour population (Cluster 1 vs Cluster 2). The top 10 terms are shown. **(E)** Category network plot showing relationships between key genes and enriched GO terms shown in **(D)**.

To assess whether this pattern reflected genuine genomic clustering rather than a random distribution of genes across the genome, we performed permutation analysis to quantify the number of upregulated gene pairs located within 2 Mb of one another following random gene shuffling (n = 5000; Figure 3B). The observed clustering of neighbouring upregulated genes was significantly greater than expected by chance (*P* = 2e-04), indicating a non-random genomic distribution of these genes, and supporting the presence of specific gene clusters associated with the inferred SCNAs. We therefore grouped neighbouring genes into gene clusters and stratified them by cluster size to identify local genomic regions underlying the inferred SCNAs (Figure 3C). This analysis highlighted particularly dense regions on 3q and 8q, coinciding with the strongest inferred arm-level gains. Consistent with the density hotspots observed in Figure 3A, several of the largest clusters were centred on established HGSOC drivers, including *MECOM*, *PIK3CA*, *MYC* and *CCND1* (Figure 3C).

Gene Ontology (GO) enrichment analysis of tumour-upregulated genes revealed strong enrichment for biological processes linked to CIN, including mitotic cell cycle progression, chromosome segregation and nuclear division (Figure 3D). Network analysis identified a coordinated gene programme involving key cancer-associated genes, such as *MKI67*, *CCNE1* and *E2F1* (Figure 3E). Together, these results link inferred chromosome arm-level copy number signal to non-random local genomic clusters containing ovarian cancer driver genes and transcriptional programmes characteristic of chromosomal instability.

### Intermediate SCNA population exhibits features of early HGSOC evolution

We next focused on Cluster 3 cells located within the histologically normal fallopian tube to investigate candidate early genomic alterations (Figure 4). These cells were concentrated along the epithelial lining of the fallopian tube (Figure 4A; left) and were strongly enriched for SECs (SECs) relative to all other cells within the normal fallopian tube region (OR = 18.3, 95% CI = 15.1-22.0, *P* < 2e-16; Figure 4A; right). Additionally, Cluster 3 was enriched for *TP53*-high cells (OR = 5.03, 95% CI = 3.97-6.47, *P* < 2e-16) (Figure 4B). Despite this enrichment, Cluster 3 cells showed reduced *TP53* target gene activity in a module comprising *CDKN1A*, *MDM2*, *GDF15*, *DDB2* and *GADD45A*, selected from high-confidence canonical *TP53* targets reported by Fischer^32^ (*P* < 2e-16; Figure 4C), with *TP53*-high cells exhibiting lower *TP53* target gene activity than *TP53*-low cells (Supplementary Figure 4). These findings are consistent with current models in which SECs and early *TP53*-associated abnormalities contribute to HGSOC initiation^9,33,34^.

**Figure 4.**
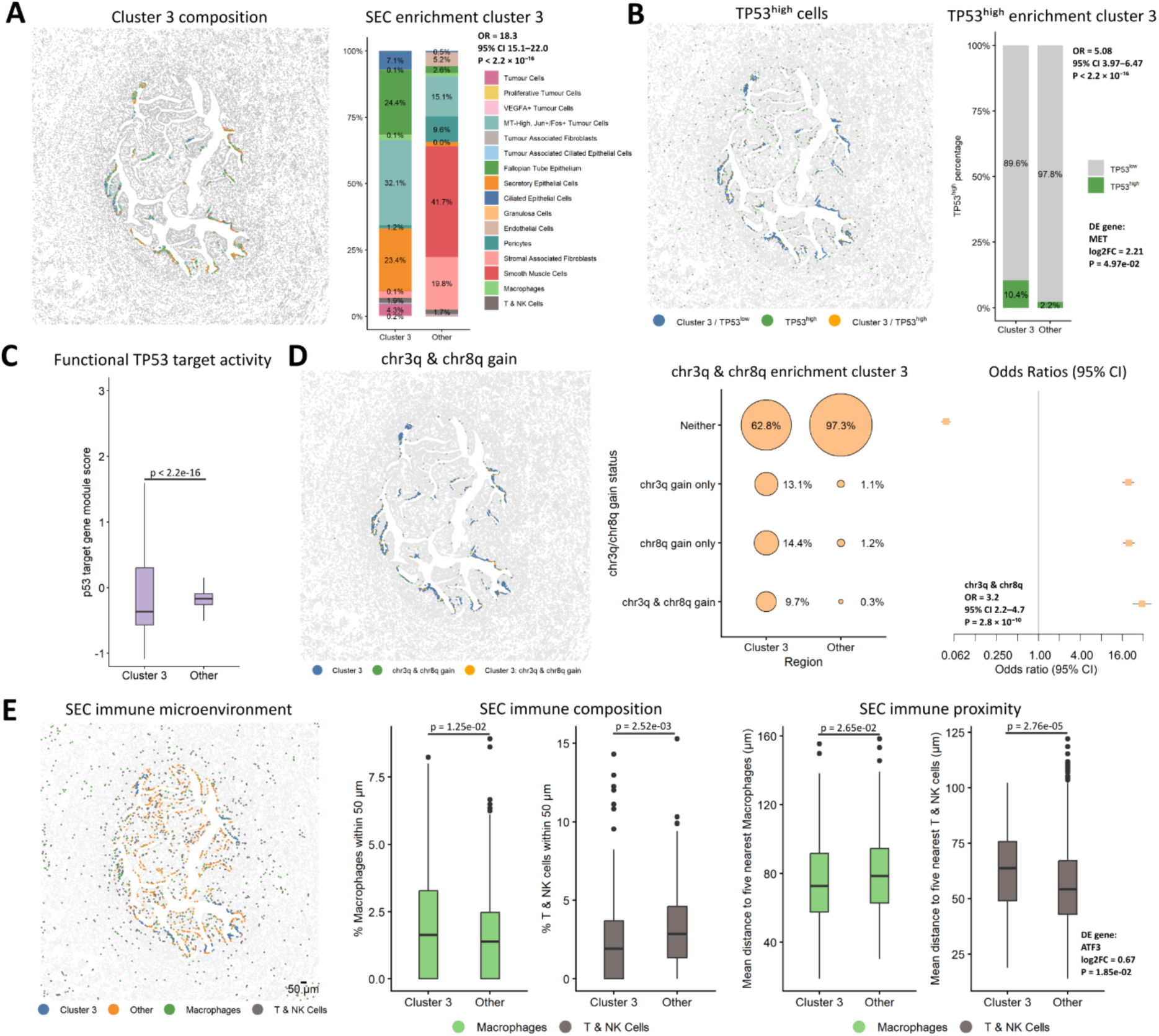
An intermediate SCNA-associated cell population within histologically normal fallopian tube epithelium is enriched for SECs (SECs), *TP53*^high^ transcriptional state, recurrent inferred 3q/8q gains and a distinct local immune architecture. (**A**) Spatial distribution of intermediate cell population (Cluster 3) within the normal fallopian tube region (left) and cell type composition of Cluster 3 compared with all other cells (right). Enrichment of SECs was assessed using Fisher’s exact test. **(B)** Spatial distribution of *TP53*^high^ cells within the normal fallopian tube region (left) and enrichment of *TP53*^high^ cells in Cluster 3 compared to all other cells in the region (right). Fisher’s exact test. *MET* was enriched in Cluster 3 (Wilcoxon rank-sum; log2FC = 2.21, *P* = 4.87e-02) (C) *TP53* target gene module score comparing Cluster 3 and other cells. Module scores were calculated from a curated set of functional *TP53* target genes defined by Fischer^32^ (Wilcoxon rank-sum test). **(D)** Spatial distribution (left) of cells harbouring both chromosome 3q gain and chromosome 8q gain within the normal fallopian tube region. Bubble plot (middle) shows the enrichment of chr3q gain, chr8q gain, and combined chromosome 3q and 8q gain in Cluster 3 compared with all other cells. Odds ratio and 95% confidence intervals (right) were estimated using Fisher’s exact test. Co-occurrence of chr3q and chr8q gains was significant (OR = 3.21, 95% CI 2.21-4.65, *P* = 2.8e-10). **(E)** Spatial distribution of immune microenvironment surrounding SECs within the normal fallopian tube region (left), stratified by Cluster 3 and outside Cluster 3 (other). Macrophage and T/NK cell abundance within 50 µm (middle), and mean distance to the five nearest macrophages and T/NK cells, were compared between groups. Significance was assessed using Wilcoxon rank-sum test with Benjamini-Hochberg multiple testing correction. *ATF3* was enriched in SECs within Cluster 3 (Wilcoxon rank-sum test; log2FC = 0.67, *P* = 1.85e-02).

Within Cluster 3, *TP53*-high cells showed increased *MET* expression relative to *TP53*-low cells (log2FC = 2.21, *P* = 0.049). As this proto-oncogene acts upstream of the PI3K pathway, its enrichment suggests a possible early dependency on pro-survival signalling that enables cells within this *TP53*-high intermediate population to tolerate CIN-associated stress. Cluster 3 was enriched for cells with inferred 3q, 8q gains, and combined 3q/8q gain relative to other cells in the normal region (Figure 4D). Notably, co-occurrence of inferred 3q and 8q gains within Cluster 3 was significant (OR = 3.21, 95% CI = 2.21-4.65, *P* = 3e-10). These data suggest that tumour-associated copy number patterns may already be detectable in a spatially restricted epithelial population before overt pathological transformation.

To assess the robustness of the intermediate population and determine whether it could be identified independently of the tumour compartment, we repeated the SCNA-based clustering analysis using only cells from the normal fallopian tube epithelium. This independent analysis largely recapitulated the intermediate cluster identified in the full dataset, with 354 of 440 intermediate cluster cells (80.5%) overlapping with Cluster 2 identified in the normal fallopian tube only analysis (Supplementary Figure 5).

We finally examined the local immune context of SECs in Cluster 3 (Figure 4E). Compared with SECs outside Cluster 3, these cells were surrounded by fewer T and natural killer (NK) cells and were located farther from neighbouring T and NK cells, whereas macrophages showed greater local abundance and closer spatial proximity (Figure 4E). Intriguingly, *ATF3* (a stress response factor implicated in tumour-associated immune remodelling^35,36^) was more highly expressed in SECs within Cluster 3 than in SECs outside Cluster 3 (log2FC = 0.67, *P* = 0.019). Together, these findings raise the hypothesis that SECs within Cluster 3 exhibit early genomic alterations accompanied by local immune remodelling, potentially promoting immune evasion and pro-tumourigenic activity^37–39^.

### SCNA inference is broadly applicable across multiple tumour types

Having demonstrated the biological relevance of inferred SCNA populations in HGSOC evolution, we next assessed the broader applicability of our copy number inference pipeline across additional tumour types using publicly available Xenium 5K datasets from human cervical and breast cancers (10x Genomics) (Figure 5). In cervical cancer, three inferred SCNA populations were identified (Figure 5A). Clusters 2 and 3 showed pronounced chromosome arm-level SCNAs, with prominent 3q, 8q and 20q gains, which are among the most frequently amplified chromosomes in cervical cancer^40,41^. In contrast, Cluster 1 remained largely copy number neutral (Figure 5A). Spatial mapping aligned these populations with distinct pathological regions (Figure 5B). Cluster 3 corresponded to in situ disease, whereas Cluster 2 mapped predominantly to invasive squamous cell carcinoma.

**Figure 5.**
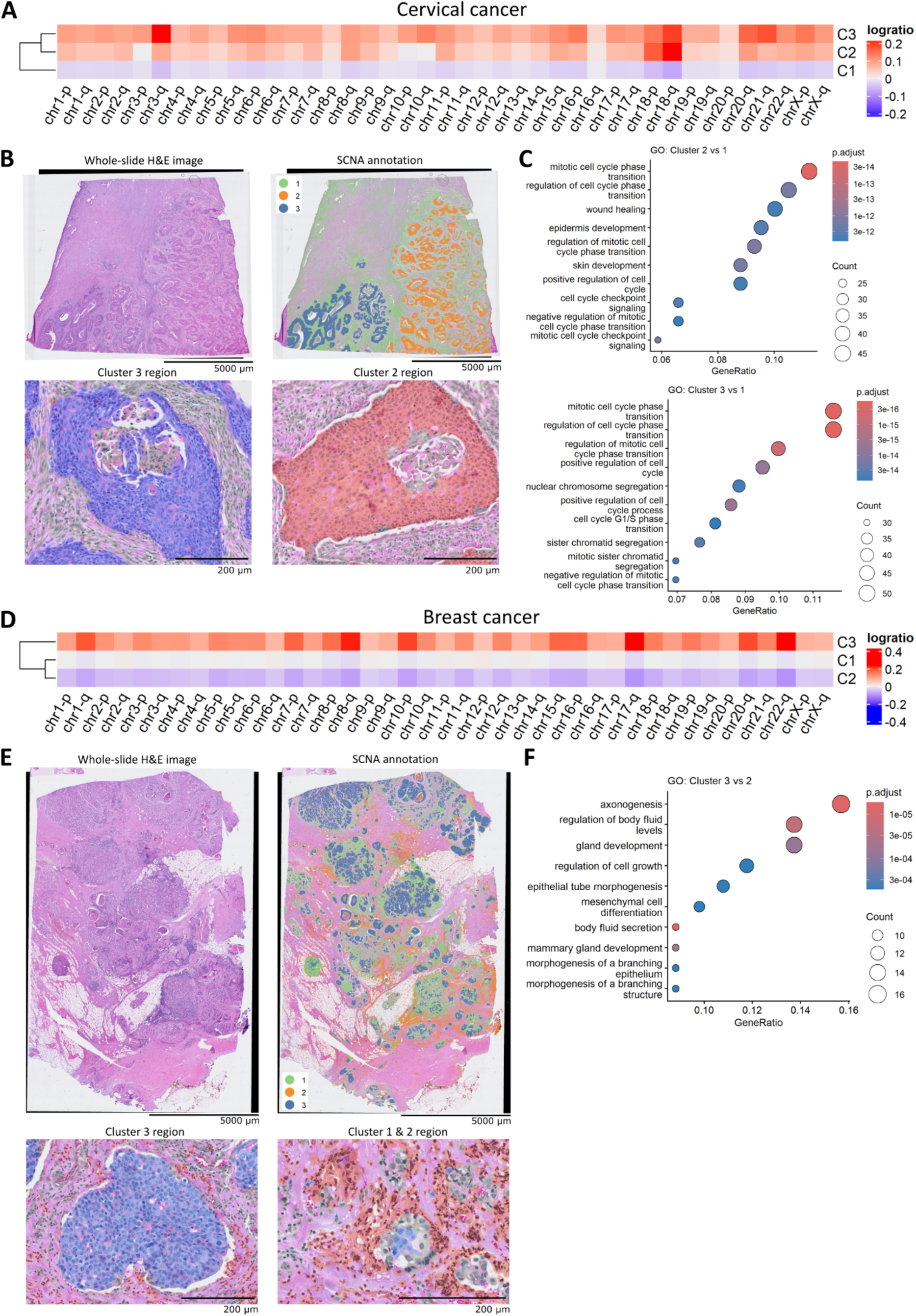
Chromosomal SCNA inference is broadly applicable across multiple tumour types. **(A)** Heatmap of spatially inferred chromosome arm-level SCNAs in Xenium 5K human cervical cancer spatial transcriptomics dataset (10x Genomics). **(B)** Whole-slide H&E image (top left), annotated SCNA clusters (top right) and representative regions corresponding to Cluster 3 and Cluster 2 (bottom). **(C)** GO enrichment analysis of genes upregulated in Cluster 2 relative to Cluster 1, and Cluster 3 relative to Cluster 1. The top 10 terms are shown. **(D)** Heatmap of spatially inferred chromosome arm-level SCNAs in Xenium 5K human breast cancer spatial transcriptomics dataset (10x Genomics). **(E)** Whole-slide H&E image (top left), SCNA cluster annotation (top right) and representative regions corresponding to Cluster 3, and Cluster 1 and 2 (bottom). **(F)** GO enrichment analysis of genes upregulated in Cluster 3 relative to Cluster 2. The top 10 terms are shown.

These findings highlight the ability of our pipeline to resolve spatial tumour populations with distinct copy number profiles associated with different pathological states, including regions containing in situ disease and invasive carcinoma. Genes upregulated in Cluster 2 and Cluster 3 relative to Cluster 1 showed strong enrichment for processes associated with chromosomal instability, including chromosome segregation, mitotic cell cycle progression and cell cycle phase transition (Figure 5C).

In the breast cancer dataset, three distinct SCNA populations were similarly resolved (Figure 5D). Cluster 3 showed the most extensive inferred chromosome arm-level SCNA gains, including at 1q, 8q and 17q, which are recurrently gained in breast cancer^3,42,43^. Accordingly, this cluster was predominantly localised to tumour regions within the tissue section (Figure 5E). In contrast, Clusters 1 and 2 remained largely copy number neutral, with Cluster 1 surrounding the Cluster 3 regions and Cluster 2 occupying more distant tissue regions. Genes upregulated in Cluster 3 relative to Cluster 2 were enriched for developmental processes including regulation of cell growth and epithelial tube morphogenesis (Figure 5F), highlighting biological processes associated with breast tumour development.

Collectively, these findings demonstrate that our copy number inference pipeline is broadly applicable across multiple tumour types, resolving biologically meaningful spatial SCNA populations associated with distinct pathological states and transcriptional programmes.

## Discussion

Here, we present a spatial transcriptomic framework that uses coordinated chromosome arm-level gene expression and spatial structure to infer broad SCNA-associated cell populations in situ. The framework identifies broad copy number aberrations, leverages spatial information to refine copy number estimates and identify subpopulations with shared copy number profiles. It further detects local hotspots of neighbouring co-expressed genes and coordinated gene clusters, and characterises the transcriptional programmes and biological pathways associated with the inferred SCNA states. In ovarian cancer, the inferred states recapitulated recurrent HGSOC-associated copy number patterns and separated tumour from non-malignant regions. Importantly, the approach identified an intermediate population within histologically normal fallopian-tube epithelium that was enriched for SECs, a *TP53*-high state and inferred 3q/8q gains. These findings are in line with the recent identification of SECs under replication stress and aneuploid cells in the normal fallopian tube epithelium^9,44^.

The spatial context of this intermediate population is particularly informative. SECs within the population showed reduced local T/NK-cell abundance and greater T/NK-cell distance, together with increased macrophage abundance and proximity. These observations connect candidate early genomic evolution with local immune organisation and raise a mechanistic hypothesis: survival of emerging genomically altered secretory clones may depend not only on cell-intrinsic tolerance of CIN but also on a microenvironment in which lymphocyte surveillance is attenuated and macrophage interactions are altered. It also suggests that the previously described immune evasion in progressive pre-cancerous lesions^45^ may occur earlier in the histologically normal fallopian tube epithelium.

Although SCNAs are inferred from transcriptional gene-dosage effects rather than directly measured DNA copy number, and there is a theoretical risk that transcriptional programmes can mimic or obscure copy number-associated expression, spatial coherence, concordance with recurrent TCGA alterations and enrichment of CIN-related programmes strengthen biological plausibility. The present analysis also establishes association, not causality, between the epithelial state and its immune context.

Applying the framework to histologically normal fallopian tubes from genetically high-risk and population-risk women, with matched single-cell DNA sequencing and detailed hormonal exposure, should establish the specificity and prevalence of these states.

Prospective human perturbation studies can then test whether preventive interventions alter the abundance, transcriptional phenotype or immune context of SCNA-associated epithelial populations. Such experiments would move the field from spatial description towards direct interrogation of mechanisms that determine whether early genomically altered clones are eliminated, persist or progress.

More broadly, the ability to map candidate genomic evolution onto intact tissue architecture provides a route to studying pre-malignant ecosystems across cancer types. By integrating spatial transcriptomics with orthogonal genomic measurements, the framework could help define how emerging clones interact with neighbouring immune and stromal cells during the earliest stages of malignant transformation.

### Limitations

Although we identify an intermediate cell population within the histologically normal fallopian tube characterised by recurrent SCNAs, TP53 dysregulation and immune remodelling, the limited sample size prevents generalisation of these observations across individuals. Future studies in larger cohorts will be required to validate the reproducibility and biological significance of these current findings.

## Methods

The main steps of the spatial SCNA inference pipeline are described below. A fully documented R package including example workflows and supporting documentation will be released upon publication in a peer-reviewed journal.

### Spatial Copy Inference pipeline Data availability and input

The following publicly available Xenium 5K datasets from 10x Genomics were used in this study: FFPE Human Ovarian Cancer with 5K Human Pan Tissue and Pathways Panel plus 100 Custom Genes (https://www.10xgenomics.com/datasets/xenium-prime-ffpe-human-ovarian-cancer), FFPE Human Breast Cancer with 5K Human Pan Tissue and Pathways Panel plus 100 Custom Genes (https://www.10xgenomics.com/datasets/xenium-prime-ffpe-human-breast-cancer) and FFPE Human Cervical Cancer with 5K Human Pan Tissue and Pathways Panel plus 100 Custom Genes (https://www.10xgenomics.com/datasets/xenium-prime-ffpe-human-cervical-cancer). The Human Ovarian Cancer FFPE Single Cell Gene Expression Flex (Next GEM) (https://www.10xgenomics.com/datasets/17k-human-ovarian-cancer-scFFPE) was used for cell type annotation.

The pipeline accepts two inputs: (i) a directory containing Xenium 5K output files and (ii) an optional cell type annotation file. Data are imported and processed as Seurat objects, ensuring compatibility with standard workflows within the Seurat toolkit^46^ .

### Quality control

Quality control was performed by removing cells with low numbers of detected genes and counts, and by filtering lowly expressed genes. Cells with numbers of detected genes and counts below the 2^nd^ percentile thresholds were removed. Genes expressed in fewer than 50 cells or with fewer than 50 total counts across the dataset were excluded from downstream analyses. All thresholds can be modified by the user.

### Normalisation

Normalisation was performed using Seurat’s *LogNormalize* method (scale.factor = 10,000), followed by conversion to log2-transformed normalised expression values.

### Chromosome-arm aggregation

Genes were mapped to chromosome arms (p and q) using hg38 genome annotations. Chromosome arms represented by fewer than five genes were removed. For each cell, chromosome arm-level expression was calculated as the mean log2-normalised expression of all genes assigned to that chromosome arm.

### Reference selection

Non-malignant cells were used as reference cells to provide a normal diploid baseline for SCNA inference. Reference chromosome-arm profiles were computed from the mean expression of the selected reference cells. When cell type annotations or an appropriate reference population are unavailable, the pipeline uses the global mean across all cells as the default reference. For the human ovarian cancer dataset, Fallopian Tube Epithelium and SECs were used as the reference population.

### SCNA inference

SCNA inference was performed by calculating chromosome arm-level log ratios. For each cell, the diploid reference baseline was subtracted from the chromosome arm-level expression. Positive and negative log ratio values indicate relative gains and losses with respect to the reference.

### Spatial clustering

To incorporate spatial information, spatial clustering was implemented using the BANKSY algorithm^27^. Chromosome arm-level log ratios (SCNAs) were used as input features, with a mixing parameter of *λ* = 0.8 and a spatial neighbourhood size of *kgeom* = 30. Principal component analysis was performed on the BANKSY-transformed chromosome arm-level log ratio profiles, and a nearest-neighbour graph was constructed using the 30 nearest neighbours across the first eight principal components. Spatial clusters were then identified using the Leiden community detection algorithm with a clustering resolution of 0.1^47^ . The clustering parameters can be modified as required for specific datasets.

### Differential expression analyses

Differential gene expression analyses were performed between spatial clusters using the Wilcoxon rank-sum test as implemented in Seurat’s *FindMarkers* function^46^. GO enrichment analyses were performed using clusterProfiler^48^ on significantly upregulated genes with thresholds of *Padj* < 0.01 and log2FC > 0.

### Gene clusters and local density

Key gene clusters underlying spatial SCNA populations were identified by grouping neighbouring genes differentially expressed between spatial clusters and located within 2 Mb of one another on the same chromosome. Gene clusters were subsequently stratified by size (2, 3, 4 or ≥ 5 genes). Local density of differentially expressed genes was estimated using overlapping 5 Mb genomic windows with a step size of 500 kb. Density scores were calculated as the number of differentially expressed genes divided by the square root of the number of panel genes within each window to account for regional differences in panel coverage, and subsequently rescaled between 0 and 1. Gene clusters and local density profiles were visualised using karyoploteR^49^.

### Normal fallopian tube region analyses

A histologically normal fallopian tube region was manually subset based on spatial coordinates and used for downstream analyses.

### Secretory epithelial cell enrichment

Cell type composition of the intermediate cell population (Cluster 3) within the normal fallopian tube was compared with all other cells in the region. Enrichment of SECs within Cluster 3 was assessed using Fisher’s exact test.

### TP53 high and target gene module score

*TP53*-high cells were defined as cells with *TP53* expression above the 10^th^ percentile among *TP53* expressing cells. Enrichment of *TP53*-high cells within Cluster 3 was assessed using Fisher’s exact test. Differential expression between *TP53*-high and *TP53*-low cells within Cluster 3 was assessed using Seurat’s *FindMarkers* function with *Padj* < 0.05. *TP53* target gene module scores were computed using Seurat’s *AddModuleScore* function from a curated set of functional *TP53* target genes. Genes were selected from the top 10 *TP53* target genes reported by Fischer^32^ that were also present in the Xenium 5K panel (*CDKN1A*, *MDM2*, *GDF15*, *DDB2* and *GADD45A*). Differences in module scores were assessed using the Wilcoxon rank-sum test for comparisons between two groups or using the Kruskal-Wallis test followed by pairwise Wilcoxon rank-sum tests with Benjamini-Hochberg multiple testing correction for comparisons involving more than two groups.

### Chromosome arm gain co-localisation

Chromosome arm-level SCNA profiles were used to investigate the co-localisation of chromosome 3q and 8q gains within the normal fallopian tube region. Cells were classified as harbouring chr3q and chr8q gains using a chromosome arm log ratio threshold of 0.1.

Enrichment of chr3q gain, chr8q gain and combined chromosome 3q/8q gain within Cluster 3 compared with all other cells within the region was assessed using Fisher’s exact tests, with effect sizes reported as odds ratios and 95% confidence intervals. Co-occurrence of chr3q and chr8q gains within Cluster 3 was additionally evaluated using Fisher’s exact test.

### Immune neighbourhood

To characterise immune remodelling within the local microenvironment, neighbourhood analyses were performed on SECs within the normal fallopian tube region. Immune neighbourhood composition was quantified as the proportion of macrophages and T/NK cells within a 50 µm radius of each SEC. Immune neighbourhood proximity was calculated as the mean distance from each SEC to its five nearest macrophages and five nearest T/NK cells.

Differences in immune neighbourhood composition and proximity were compared between SECs within Cluster 3 and outside Cluster 3 using Wilcoxon rank-sum tests with Benjamini-Hochberg multiple testing correction. Differential expression between SECs within Cluster 3 and outside Cluster 3 was assessed using Seurat’s *FindMarkers* function with *Padj* < 0.05.

## Acknowledgements

This work was supported by the Cancer Research UK International Alliance for Cancer Early Detection Programme Grant: ACEPGM-2023/100001. ACA is supported by Cancer Research UK grant: SEBCD3-2024/100001. F.C.M. is supported by the UCL Institute for Women’s Health and Cancer Institute. This work was also supported by core funding from the NIHR UCLH Biomedical Research Centre (BRC1113/CN/FM/101330) and the NIHR Cambridge Biomedical Research Centre (NIHR203312) [*]. *The views expressed are those of the author(s) and not necessarily those of the NIHR or the Department of Health and Social Care.

## Supplementary Information

**Supplementary Figure 1.**
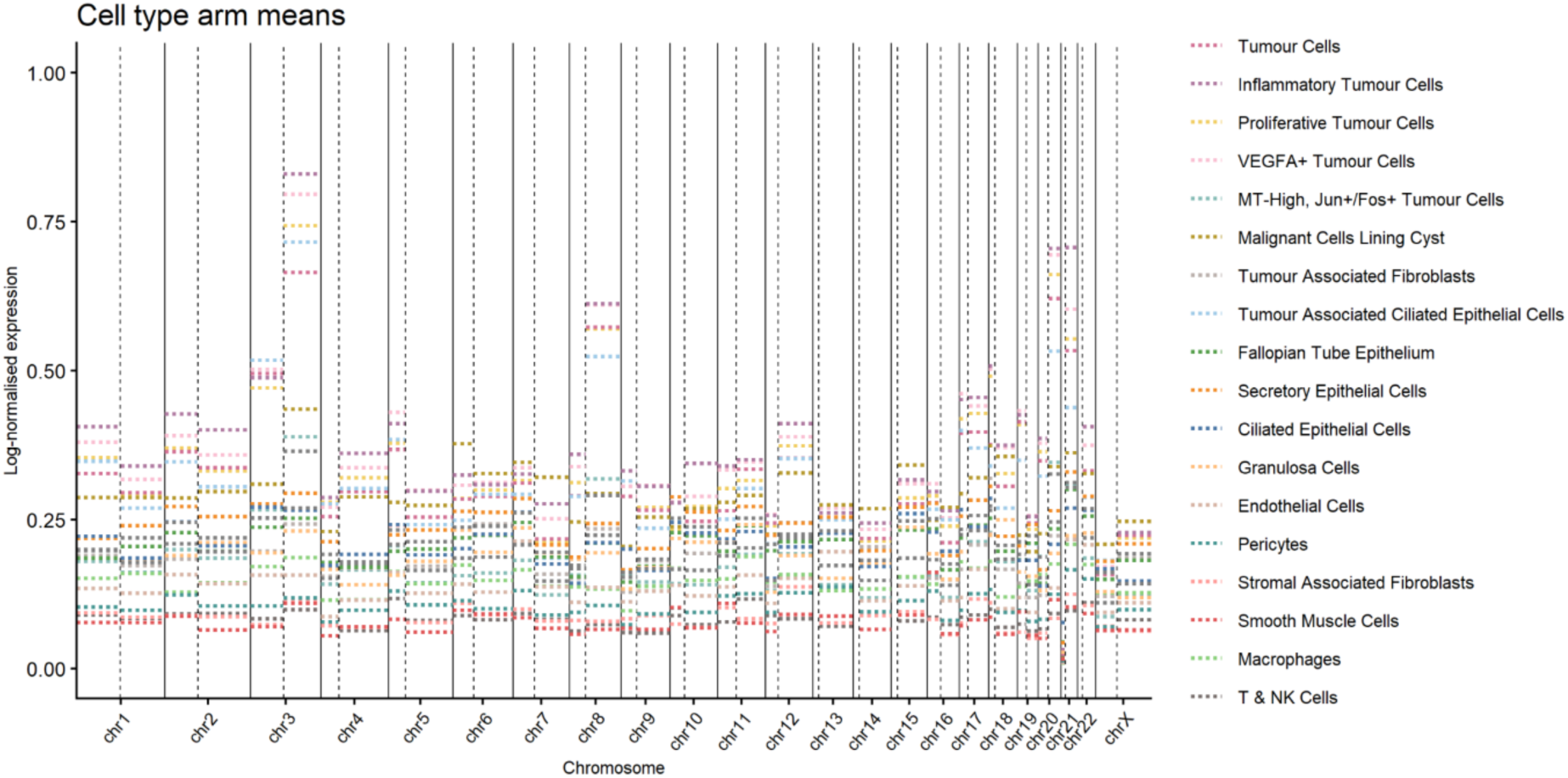
Mean chromosome arm-level gene expression across different cell types.

**Supplementary Figure 2.**
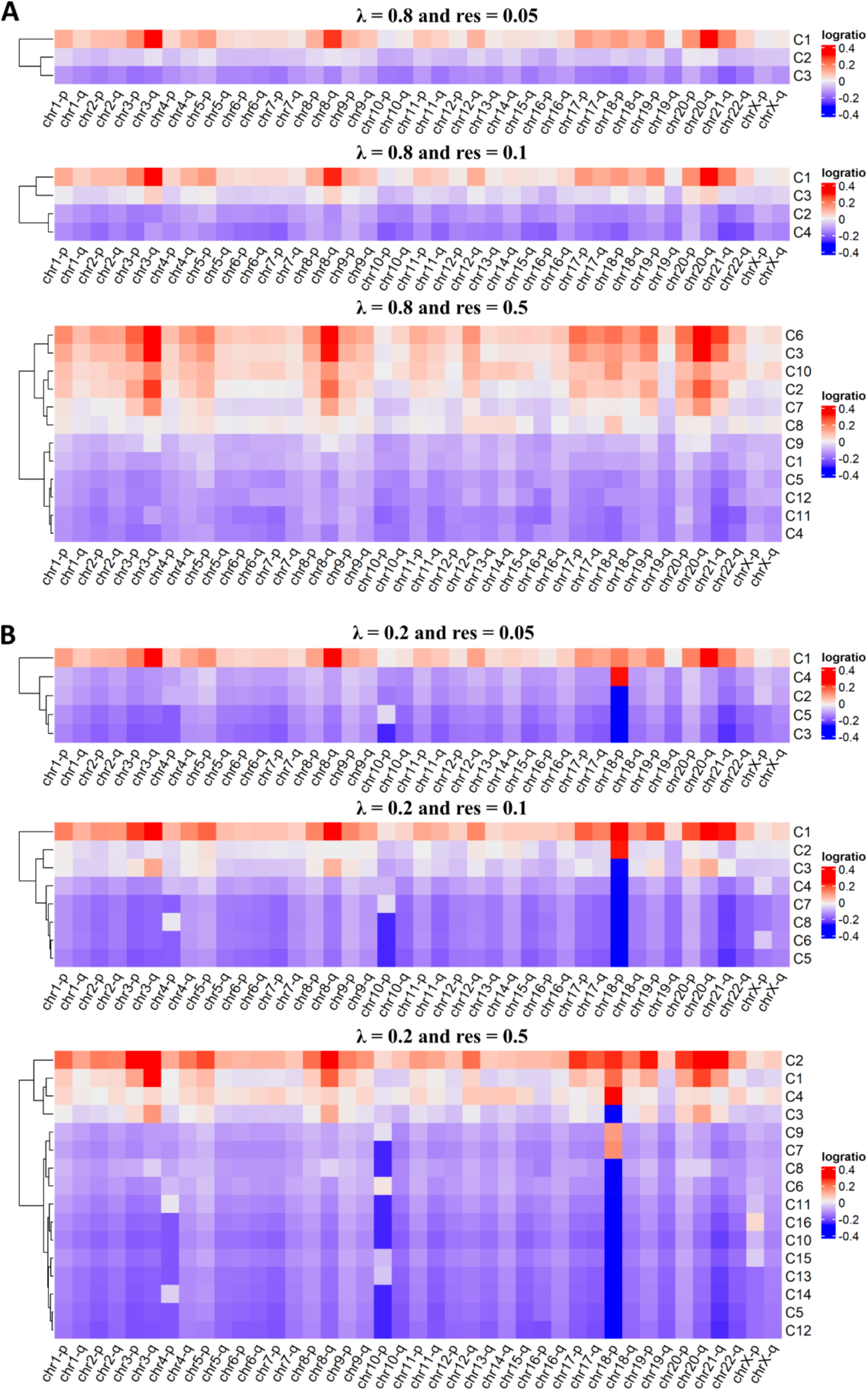
Heatmaps of chromosome arm-level SCNA log ratios across clusters identified at different values of the mixing parameter (*λ*) and clustering resolution (res). **(A)** *λ* = 0.8; res = 0.05, 0.1 and 0.5. **(B)** *λ* = 0.2; res = 0.05, 0.1 and 0.5.

**Supplementary Figure 3.**
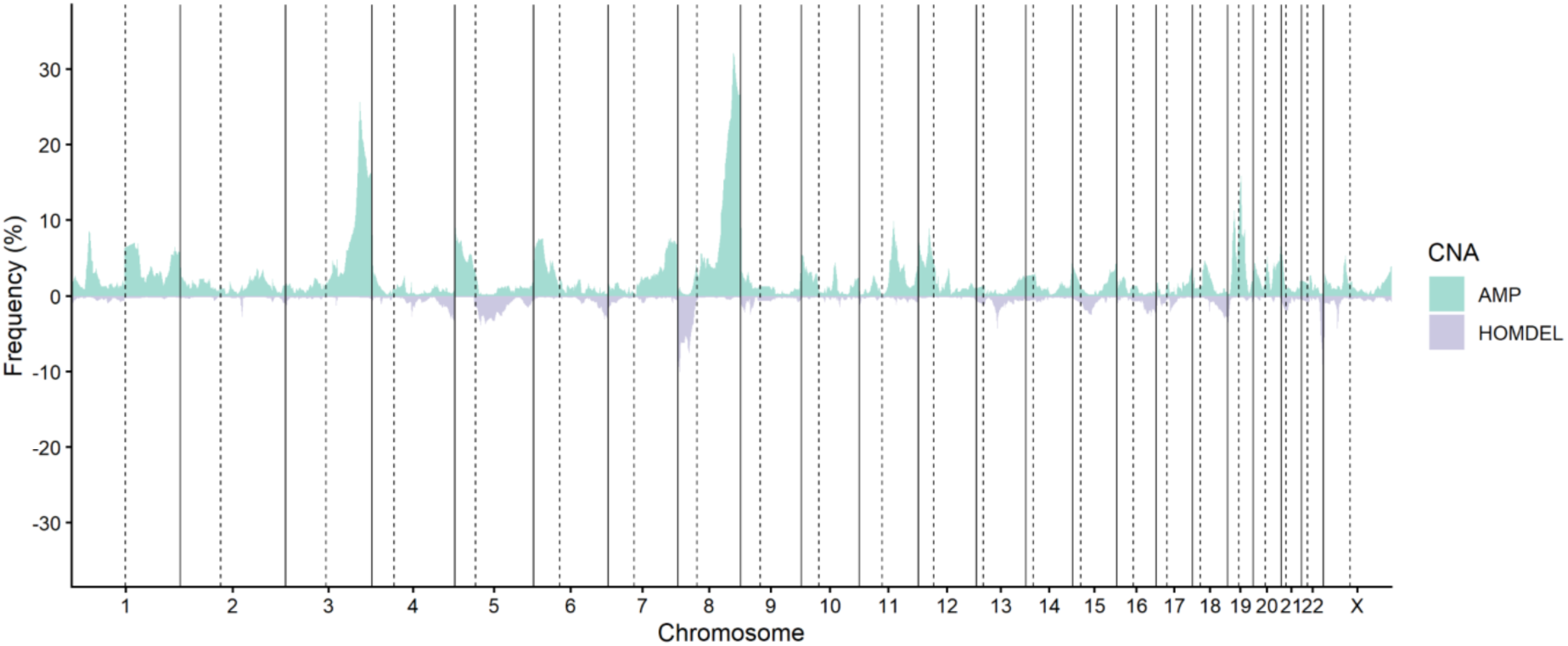
Genome-wide frequency of somatic copy number alterations in TCGA HGSOC cohort. Frequencies of copy number amplifications (AMP; green) and homozygous deletions (HOMDEL; purple) were obtained from the TCGA Ovarian Serous Cystadenocarcinoma PanCancer Atlas cohort through cBioPortal (n=572)^4,28,29^.

**Supplementary Figure 4.**
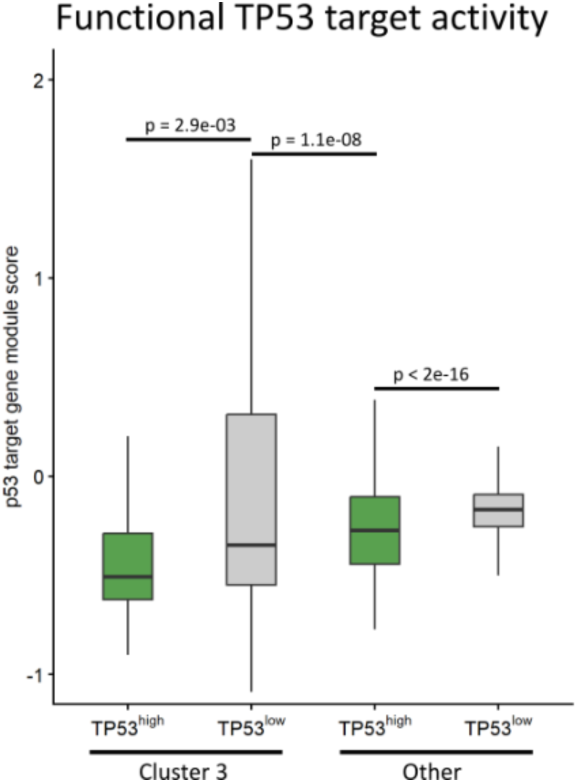
*TP53* target gene module scores comparing Cluster 3 and other cells stratified by *TP53*-high and *TP53*-low groups. Module scores were calculated from a curated set of functional *TP53* target genes defined by Fischer^32^ . Significance was assessed using a Kruskal-Wallis test followed by post hoc pairwise Wilcoxon rank-sum tests with Benjamini-Hochberg multiple testing correction.

**Supplementary Figure 5.**
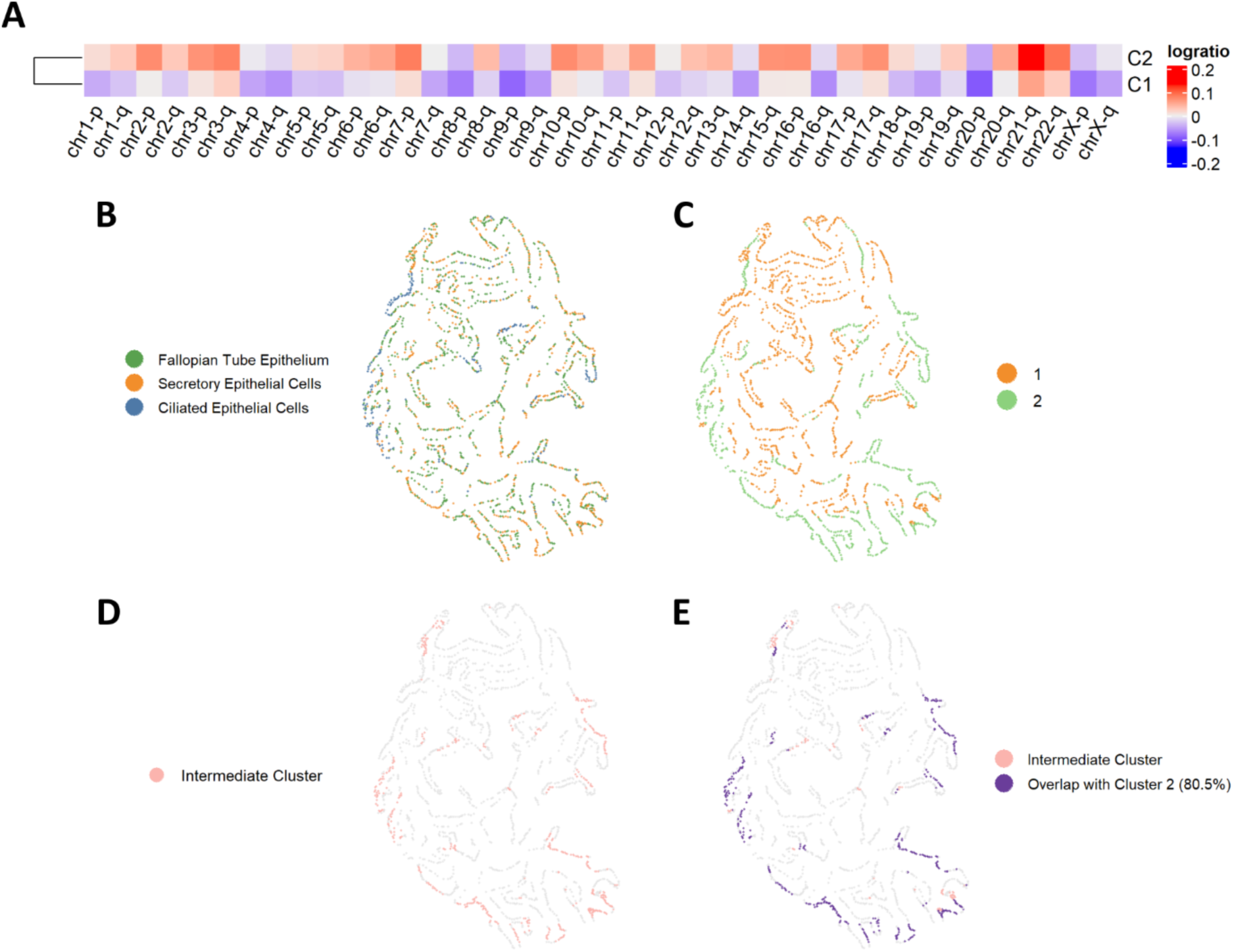
Independent SCNA-based clustering of the normal fallopian tube epithelium largely recapitulates the intermediate cluster identified in the full population. **(A)** Heatmap of chromosome arm-level SCNA log ratio profiles across clusters identified in the normal fallopian tube epithelium population. **(B)** Cell type distribution of the normal fallopian tube epithelium population used for SCNA-based clustering. Colours indicate cell types. **(C)** Spatial distribution of the SCNA-defined clusters identified within the normal fallopian tube epithelium population. Colours indicate SCNA-defined clusters. **(D)** Spatial distribution of the intermediate cluster identified in the full population within the normal fallopian tube epithelium. **(E)** Overlap between the intermediate cluster and SCNA-defined Cluster 2. A total of 354 of 440 intermediate cluster cells (80.5%) overlapped with SCNA-defined Cluster 2.

